# Joint genetic analysis using variant sets reveals polygenic gene-context interactions

**DOI:** 10.1101/097477

**Authors:** Francesco Paolo Casale, Danilo Horta, Barbara Rakitsch, Oliver Stegle

## Abstract

Joint genetic models for multiple traits have helped to enhance association analyses. Most existing multi-trait models have been designed to increase power for detecting associations, whereas the analysis of interactions has received considerably less attention. Here, we propose iSet, a method based on linear mixed models to test for interactions between sets of variants and environmental states or other contexts. Our model generalizes previous interaction tests and in particular provides a test for local differences in the genetic architecture between contexts. We first use simulations to validate iSet before applying the model to the analysis of genotype-environment interactions in an eQTL study. Our model retrieves a larger number of interactions than alternative methods and reveals that up to 20% of cases show context-specific configurations of causal variants. Finally, we apply iSet to test for sub-group specific genetic effects in human lipid levels in a large human cohort, where we identify a gene-sex interaction for C-reactive protein that is missed by alternative methods.

**Author summary:** Genetic effects on phenotypes can depend on external contexts, including environment. Statistical tests for identifying such interactions are important to understand how individual genetic variants may act in different contexts. Interaction effects can either be studied using measurements of a given phenotype in different contexts, under the same genetic backgrounds, or by stratifying a population into subgroups. Here, we derive a method based on linear mixed models that can be applied to both of these designs. iSet enables testing for interactions between context and sets of variants, and accounts for polygenic effects. We validate our model using simulations, before applying it to the genetic analysis of gene expression studies and genome-wide association studies of human blood lipid levels. We find that modeling interactions with variant sets offers increased power, thereby uncovering interactions that cannot be detected by alternative methods.

## Introduction

Understanding genetic interactions with external context (GxC), including environment, is a major challenge in quantitative genetics. Linear mixed models (LMMs) have emerged as the framework of choice for many genetic analyses, mainly because the random effect component in this class of models provides robust control for population structure [1, 2] and other confounding factors [3-5]. More recently, random-effect models have also been shown to be effective to test for polygenic effects from multiple causal variants that are in linkage [6–9] (variant sets). Additionally, multivariate formulations of LMMs have been developed to test for genetic effects across multiple correlated traits, both in single-variant analyses [10, 11] and more recently for joint tests using variant sets [12]. However, these existing multivariate LMMs have primarily been designed to increase the statistical power for detecting association signals, whereas methods to test for interactions are only beginning to emerge [10, 13].

Classical single-variant models for GxC use fixed effects to test for differential effect sizes of individual variants between contexts, either using an ANOVA [14–16] or LMMs [10, 17]. The main advantages of set-based tests compared to single-variant models are twofold. First, set tests reduce the effective number of tests and can account for effects due to multiple causal variants, thus increasing power for detecting polygenic effects [7, 8, 12, 18]. Second, we here show that joint tests across multiple contexts and sets of variants allow for characterizing the local architecture of polygenic-GxC interactions.

One way to generalize single-variant interaction tests to variant sets is using a model that assumes that context differences cause the same fold-differences in effect size across all genetic variants, such that all genetic effects in one context are proportional to the effects in a second context; a criterion that has also been considered to assess co-localization of multiple traits [19] (Fig. 1a, middle). We denote this class of interactions *rescaling*-*GxC*. More generally, however, there may also be differences in the configuration of causal variants between contexts (Fig. 1a, right), such that not all genetic variants show the same fold-difference between contexts, as some become more prominent in particular contexts and others less so. We denote these complex interactions *heterogeneity*-*GxC*. These two classes of interactions have different functional implications – the former suggest no difference in causal variants between contexts, and the latter suggest otherwise. Distinguishing between them is only possible using multi-variant models such as set tests, and is important for identifying different potential causal variants in different contexts.

**Figure 1.**
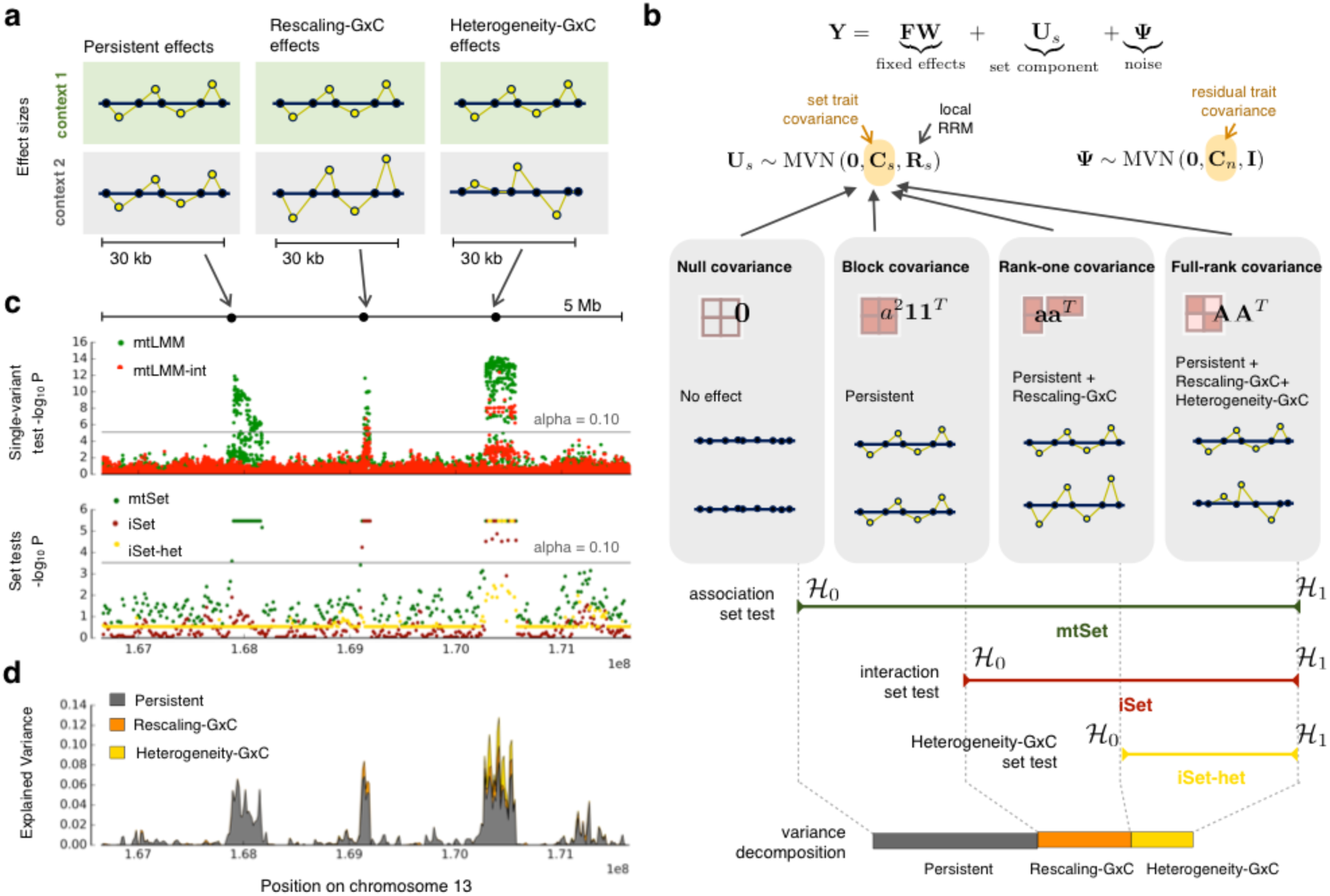
Illustration of the iSet model and different architectures of genotype-context interactions. **(a)** Alternative genetic architectures that are explicitly modeled in iSet: *persistent effects,* where causal variants have identical effects across contexts (left panel), *rescaling*-*GxC effects*, where the effects of causal variants in one context are proportional to those in a second contexts (middle), and *heterogeneity*-*GxC effects*, with changes of causal variants or their relative effect sizes between contexts (right). **(b)** Illustration of the multivariate linear mixed model (LMM) that underlies iSet. Model comparisons of LMMs with different trait-context covariance of the set component ***C****_s_* are used to define tests for general associations (*mtSet*), interactions (*iSet*) and heterogeneity-GxC effects (*iSet*-*het*). Additionally, the model can be used to estimate the proportion of variance that can be attributed to the corresponding genetic architectures **(Methods). (c,d)** Applications of iSet to a small simulated region. The total genetic effect was simulated as the sum of contributions from three loci with a persistent (left), rescaling-GxC (middle) and heterogeneity-GxC effects (right). **(c)** Manhattan plots of P values from a single-variant LMM [10] to test for associations (mtLMM) or interactions (mtLMM-int). Lower panel: Corresponding Manhattan plots for P values from set tests, considering a test for associations (mtSet), interactions (iSet) or heterogeneity-GxC (iSet-het), using consecutive regions (30 kb regions; step size 15 kb). Horizontal lines correspond to the *α* = 0.10 significance threshold (Bonferroni adjusted). P values of set tests are bounded (>10^-6^) by the number null model simulations to estimate significance levels (*Methods*). (d) Proportion of variance attributable to persistent effects, rescaling-GxC and heterogeneity-GxC, considering the same regions as in **c**.

We here propose a multivariate LMM to test for interactions test between <u>**Sets**</u> of genetic variants and categorical contexts (iSet) and to distinguish between *rescaling*-*GxC* and *heterogeneity*-*GxC*. We find that iSet yields increased power for identifying interactions and uniquely is able to robustly differentiate between rescaling-GxC and heterogeneity-GxC. We first validate iSet using simulations before applying the model to test for gene-by-sex interactions in blood lipid levels [20] as well as gene-by-environment interactions in an expression quantitative trait loci (eQTL) study [21]. We identify up to 20% of the stimulus-specific eQTLs as cases of *heterogeneity*-*GxC*, suggesting that context-specific causal variants are common.

## Results

### A mixed model approach to test for polygenic GxC

iSet generalizes previous multi-trait set tests [12], while considering the same trait measured in two (environmental) contexts. For a fully observed design, where the trait is measured in *N* individuals and each context, the phenotype matrix ***Y*** is modeled as the sum of a genetic effect from a set component and residual noise:

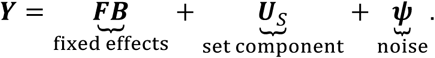

Here, ***F*** and ***B*** denote the design and the effect size matrices of additional fixed effect covariates and ***U****_s_* and ***ψ*** are random effects that follow matrix-variate normal distributions:

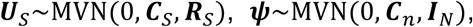

Where ***R****_S_* corresponds to a local realized relatedness matrix [22] of the set of interest *s*, and ***I**_N_* denotes a diagonal covariance, which corresponds to independent and Identically distributed residuals. The trait-context covariance matrices ***C****_s_* and ***C****_n_* model correlations between contexts due to the set component (***C****_s_*), and residual noise (***C****_n_*).

A key insight derived here is that different assumptions on the structure of the trait-context covariance ***C****_s_* correspond to alternative genetic architectures that can be explained by a polygenic model (Fig. 1b, **Methods**). *Persistent* genetic effects across contexts (no GxC) can be modeled using an LMM with a constant block covariance (Fig. 1b); *rescaling GxC*, where effect sizes in different contexts are proportional to each other, can be captured by a trait-covariance with rank one. Note that genetic effects that act only in one context are a special case of this model and corresponds to a zero rescaling coefficient. Finally, the most general architectures with different relative effect sizes between contexts (*heterogeneity*-*GxC*) can be captured by an LMM with a full-rank trait-context covariance (**Methods**). By comparing LMMs with these alternative covariance structures, it is possible to define set tests for general associations (*mtSet*), which identifies both persistent and context-specific effects, a test for genetic interactions, both with or without changes in the configuration of causal variants (*iSet*), and finally a test for heterogeneity-GxC effects (*iSet*-*het*), which is specific to differences between contexts that cannot be explained by rescaling (Fig. 1b).

These multivariate LMMs can be fit using principles we have previously derived for multivariate set tests [12], and hence iSet can be applied to large cohorts with up to one hundred thousand individuals (**Supplementary Fig. 1**). Permutation schemes are not well defined for interaction models [23], so we use a parametric bootstrap procedure [23] to estimate P values. An important advantage compared to previous interaction tests [13, 24–28] (**Methods**), is that iSet can be applied both to study designs where all individuals have been phenotyped in each context and when stratifying populations into distinct subgroups using a context variable **(Supplementary Fig. 2**). iSet also provides control for population structure, either using principal components that are included as fixed covariates, or using an additional random effect term (**Methods**). Finally, iSet can also be used to estimate the total phenotypic variance explained by variant sets and the relative proportions captured by persistent, rescaling-GxC and heterogeneity-GxC effects (**Methods**).

To illustrate the polygenic interactions that can be detected using iSet, we first considered a basic simulated example (Fig. 1c). We simulated genetic effects for one quantitative trait in two contexts, considering polygenic effects at three distinct loci (**Methods**): a region with persistent genetic effects, a region with rescaling-GxC and a region with context-specific causal variants (heterogeneity-GxC effects). We tested consecutive regions (30kb region, 15kb step) using the three tests provided by our model (mtSet, iSet, iSet-het), finding that by combining these results, it was indeed possible to resolve the architecture of each of the simulated regions (Fig. 1c-d). In particular, this example illustrates that, unlike single-variant tests, iSet-het can be used to discern heterogeneity-GxC effects specifically.

### Simulated data

Next, we used simulations based on genotypes from the 1000 Genomes project [29] to assess the statistical calibration and power of iSet. We generated a population of 1,000 individuals based on genotype data from European populations, initially simulating one quantitative trait measured in two distinct contexts in all individuals (**Methods**).

First, we considered data with simulated persistent polygenic effects, confirming that both iSet and iSet-het are calibrated when no interaction effects are simulated (Fig. 2a, **Supplementary Table 1**). Analogously, we also confirmed that iSet-het is calibrated when only rescaling-GxC effects are considered (**Supplementary Fig. 3**), and we assessed the robustness of iSet to different types of model misspecification (**Supplementary Fig. 4, Supplementary Table 1**).

**Figure 2.**
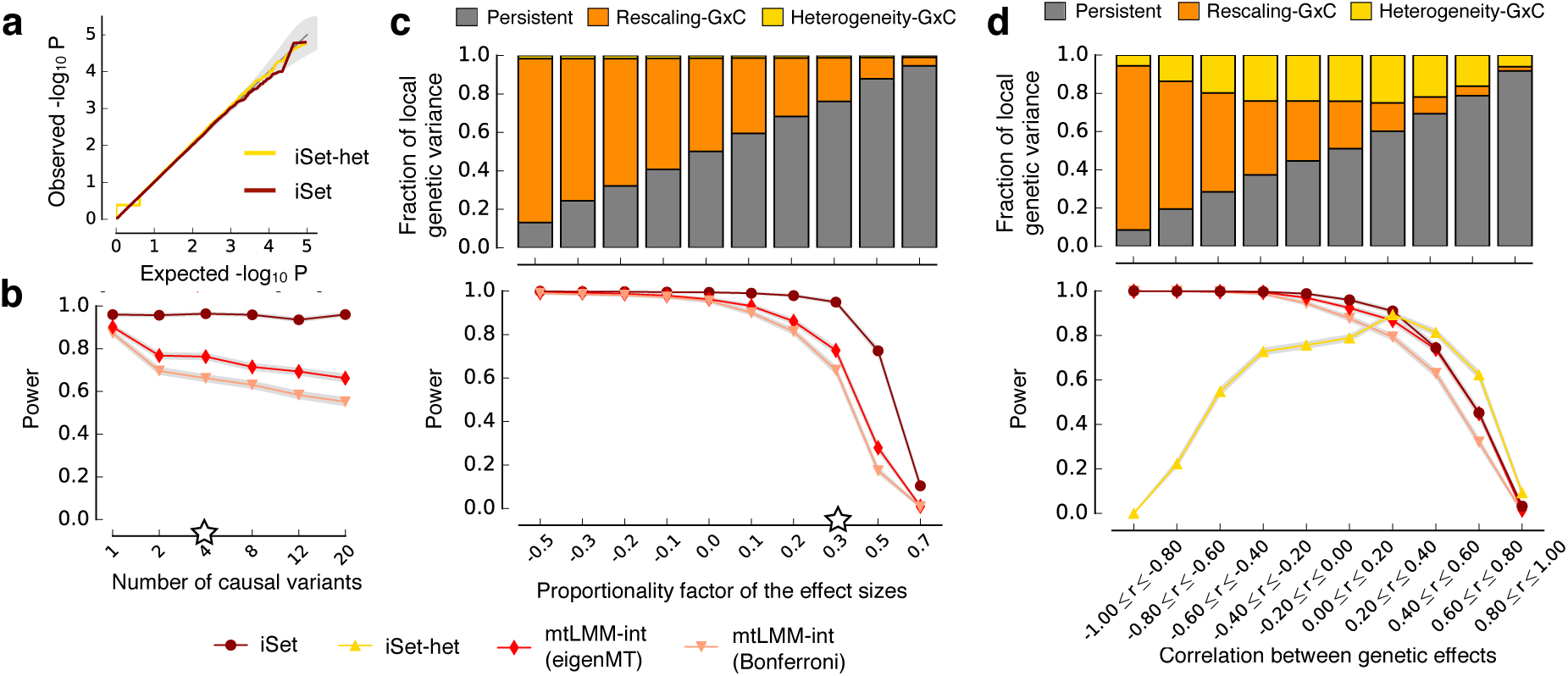
Simulated data to assess statistical calibration and power of iSet. **(a)** QQ plot for the P values obtained from iSet and iSet-het when only persistent genetic effects were simulated. The step in the QQ-plot for large p-values is observed because the trait-context covariances are required to be positive-semidefinite. **(b)** Power comparison of alternative models for detecting simulated interactions, considering rescaling-GxC effects (without heterogeneity-GxC) for increasing numbers of simulated causal variants. Compared were iSet and a single-variant interaction test (mtLMM-int) [10], using two alternative approaches to adjust for multiple testing of single variant methods (Bonferroni or eigenMT). **(c)** Lower panel: analogous power comparison as in **b**, when varying the proportionality factor of effect sizes between contexts. A proportionality factor of zero corresponds to genetic effects that act only in one of the contexts. See **Supplementary Table 3** for the relationship of the proportionality factor and fold differences. iSet-het was not considered, because all simulated rescaling-GxC are consistent with the null model of iSet-het. Top panel: average fraction of genetic variance attributable to persistent, rescaling-GxC and heterogeneity-GxC effects for the corresponding simulations. **(d)** Analogous comparison as in **c** but for simulated heterogeneity-GxC effects, when varying the correlation of the total genetic effect between contexts. Additionally, we also considered iSet-het to test for heterogeneity-GxC, which was best powered for heterogeneity-GxC effects that were uncorrelated between contexts. White stars denote default parameter values that were kept constant when varying other parameters (**Supplementary Table 2**). Statistical power was assessed at 5% FDR across 1,000 repeat experiments.

We compared iSet to single-variant interaction tests [10] (mtLMM-int) (**Methods**), considering a wide range of different settings (**Supplementary Table 2**, Methods). Because single-variant methods perform one test for each variant in the set (**Supplementary Fig. 5**), we adjusted for multiple testing using one of two approaches: i) conservative Bonferroni adjustment (Bonferroni) or ii) a recently proposed method that estimates the effective number of independent tests based on the local structure of linkage disequilibrium (LD) [30] (eigenMT). Note that existing set-based interaction tests cannot be applied to complete designs with repeat measurements and hence were not considered (**Methods, Supplementary methods**); see below and Fig. 5 for additional experiments where these methods were used. As expected, the power advantages of iSet compared to single-variant models were largest when multiple causal variants were simulated (Fig. 2b). However, iSet was better powered than mtLMM-int even for a single causal variant. Identical simulations based on synthetic independent genotypes (**Supplementary Fig. 6**) revealed that this effect is predominantly due to local LD and advantages due the reduced number of total tests. We also considered the impact of different proportionality factors of genetic effects between contexts. All models were best powered to detect GxC for negative proportionality factors (opposite effects), or when the proportionality factor was close to zero (context-specific effects) (Fig. 2c).

Next, we simulated traits with context-specific causal variants (heterogeneity-GxC). Heterogeneity-GxC is detectable when there is a change in the local causal configuration, which corresponds to the absolute correlation of local genetic effects between contexts (*r*) smaller than 1; the greater the heterogeneity GxC effects, the smaller the absolute correlation. Presence of GxC effects under tightly correlated genetic effects (*r* ≈ ±1) cannot be distinguished from rescaling-GxC. To simulate these different settings, we randomly selected two causal variants in each context and varied the extent of correlations of the genetic effect between contexts (**Fig. 2d**). When using iSet-het for detecting heterogeneity-GxC effects, the model was best powered when there is a moderate to large change in causal configuration, corresponding to low correlated genetic effects (>70% power for r^2^ < 0.16, Fig. 2d). We also considered additional settings with larger numbers of causal variants (**Supplementary Fig. 8**), and we assessed the accuracy of iSet-het to classify interaction effects into heterogeneity-GxC or rescaling-GxC effects (**Supplementary Fig. 7**, Methods). Taken together, these results confirm that iSet-het is a robust test for heterogeneity-GxC.

We also investigated the proportion of local genetic variance that can be explained by models with persistent, rescaling-GxC and heterogeneity-GxC for the corresponding simulations (Fig. 2c-d, **Methods**). The persistent effect model explained large proportions of the simulated genetic variance, even in the presence of positively correlated GxC, but could not capture variance due to GxC effects with negative rescaling (Fig. 2c,d). An LMM that models rescaling-GxC did account for negative and positive rescaling, and captured some of the heterogeneity-GxC effects (Fig. 2d). Finally, variance contributions that were exclusively captured by a heterogeneity-GxC model were largest for uncorrelated context-specific genetic effects, the same regime where the corresponding test is best powered (Fig. 2d). We also confirmed that the most flexible heterogeneity-GxC model yields unbiased estimates of the total genetic variance in genomic regions, whereas other models were biased for some simulated architectures (**Supplementary Fig. 9**).

Finally, we considered simulations where we varied both the size of the testing region and the simulated causal region, using a sliding window analysis (**Supplementary Fig. 10, Methods**). Overall, iSet was markedly robust to the window size, and was best powered when the sizes of the testing approached the size of the simulated causal region, which is in line with previous findings for set-based association testing [12]. We also observed that iSet-het is best powered for small causal regions (up to 100kb), and the power for detecting heterogeneity-GxC deteriorated when analyzing larger regions.

### Analysis of stimulus-specific eQTLs in monocytes

We next applied iSet to test for stimulus-specific genetic effects in a monocyte stimulus eQTL study [21]. We considered gene expression profiles for 228 individuals in four stimulus contexts: naive state (no stimulation), 24 hours after stimulation with interferon-γ (IFN), and stimulation with lipopolysaccharide (LPS) after two and 24 hours.

We applied iSet to test for pairwise interaction effects, considering the naive monocyte state and each stimulus condition in turn, performing a single test using proximal *cis* acting variants (plus or minus 50kb from the transcription start site; **Methods**). After quality control, we considered 12,677 probes and tested for *cis* associations (mtSet), GxC interactions (iSet) and for heterogeneity-GxC effects (iSet-het). For comparison, we also considered a conventional multi-trait LMM [10] and tested for associations and interactions in the same genomic regions, using eigenMT [30] to adjust for multiple testing (**Methods**). Although there was substantial overlap of the probes and stimulus conditions for which different methods identified significant interactions (Fig. 3b), iSet was better powered (32.7% power increase; 5,068 versus 3,818 probes and stimuli with an interaction; FDR<5%, Fig. 3a, **Supplementary Fig. 11-12, Supplementary table 4**). Additionally, iSet-het identified 1,135 probes and stimulus contexts with significant heterogeneity-GxC effects (Fig. 3a,b). This shows that a substantial proportion of stimulus-specific eQTLs are associated differences in the configuration of causal variants, suggesting context-specific regulatory architectures.

**Figure 3.**
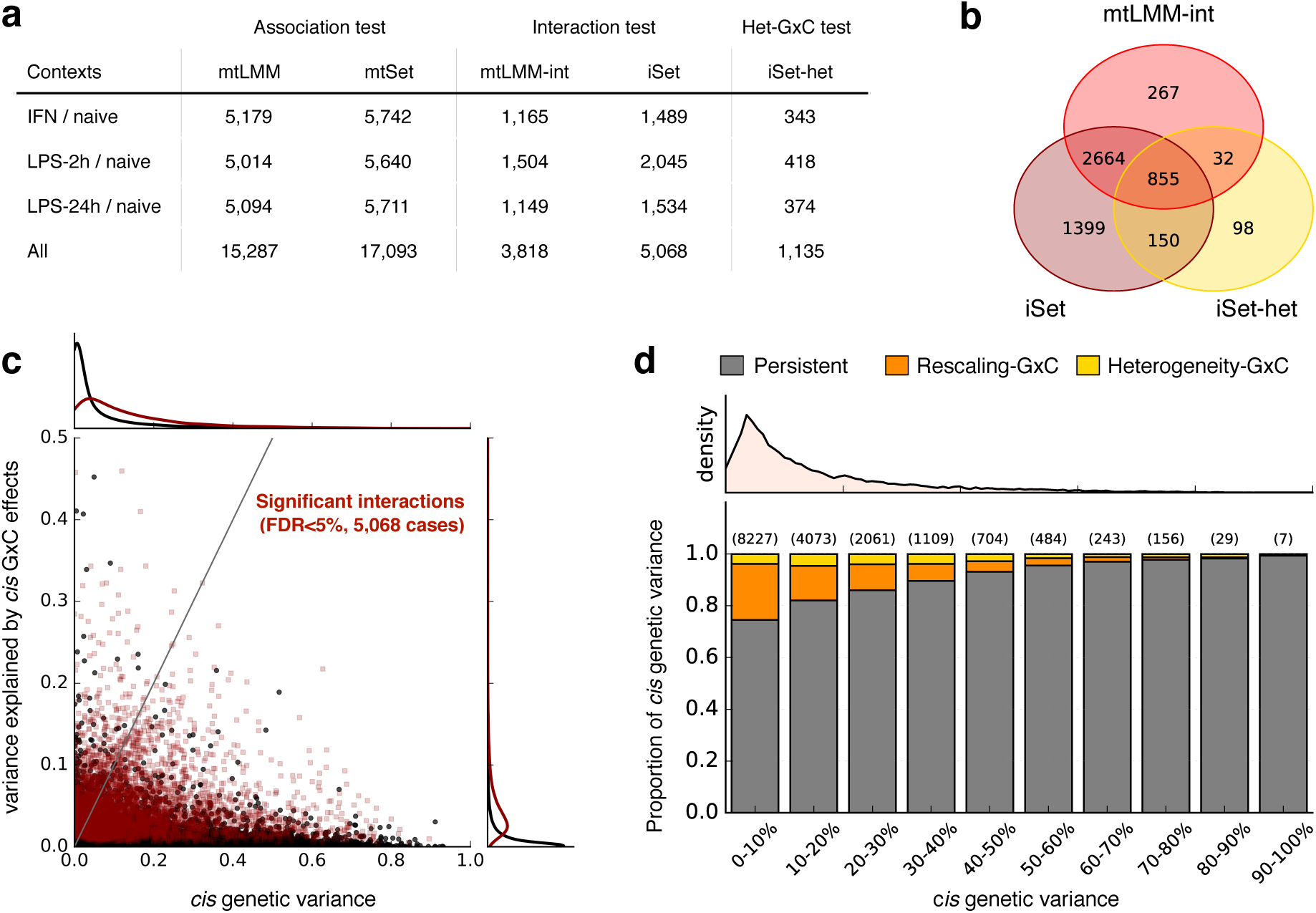
Analysis of stimulus-specific eQTLs in monocytes. **(a)** Number of probes with at least one significant *cis* association (Association test) or genotype-stimulus interaction (Interaction test) for alternative methods and stimulus contexts. Considered were the proposed set tests (mtSet, iSet, iSet-het) as well as single-variant multi-trait LMMs (mtLMM, mtLMM-int [10]), testing for genetic effects in *cis* (100kb region centered on the transcription start site; FDR < 5%). Additionally, iSet-het was used to test for heterogeneity-GxC effects. Individual rows correspond to different stimulus contexts with “All” denoting the total number of significant effects across all stimulus contexts. **(b)** Venn diagram of probes and stimuli with significant interactions identified by alternative methods and tests (across all stimuli). **(c)** Bivariate plot of the variance attributed to persistent genetic effects versus genotype-stimulus interactions for all probes and stimuli. Significant interactions are shown in red. Density plots along the axes indicate the marginal distributions of persistent genetic variance (top) and variance due to interaction effects (right), either considering all (black) or probe/stimulus pairs with significant interactions (iSet in **a**, dark red). **(d)** Average proportions of *cis* genetic variance attributable to persistent effects, rescaling effects and heterogeneity-GxC, considering probe/stimulus pairs with significant *cis* effects (5% FDR, mtSet), stratified by increasing fractions of the total *cis* genetic variance. Shown on top of each bar is the number of instances in each variance bin. The top panel shows the density of probes as a function of the total *cis* genetic variance.

Although on average the proportion of variance explained by GxC tended to be smaller than for persistent effects (median 3.7% for GxC versus median 9.5% for persistent effects, for probes with significant GxC, Fig. 3c), GxC was the driving genetic source of variation for 11.8% of the significant *cis* eQTLs (Fig. 3d; defined as explaining 50% or more of the *cis* genetic variance). Consistent with previous reports [31, 32], we observed that genes with large relative GxC effects were associated with weak overall *cis* effects, whereas eQTLs with large effect sizes tended to be persistent across stimuli (Fig. 3d).

### Mechanistic underpinning of heterogeneity eQTLs

To better understand the mechanisms that underlie genes with detected heterogeneity-GxC effects, we used an LMM with step-wise selection [33], identifying 15,756, 2,690 and 457 eQTLs (across all probes and contexts) with a single significant association, significant secondary and significant tertiary associations respectively (FDR < 5%, **Methods, Supplementary Table 4**). Probes with significant heterogeneity-GxC were more likely to harbor multiple independent associations (Fig. 4a), confirming that heterogeneity-GxC eQTLs have complex genetic architectures.

**Figure 4.**
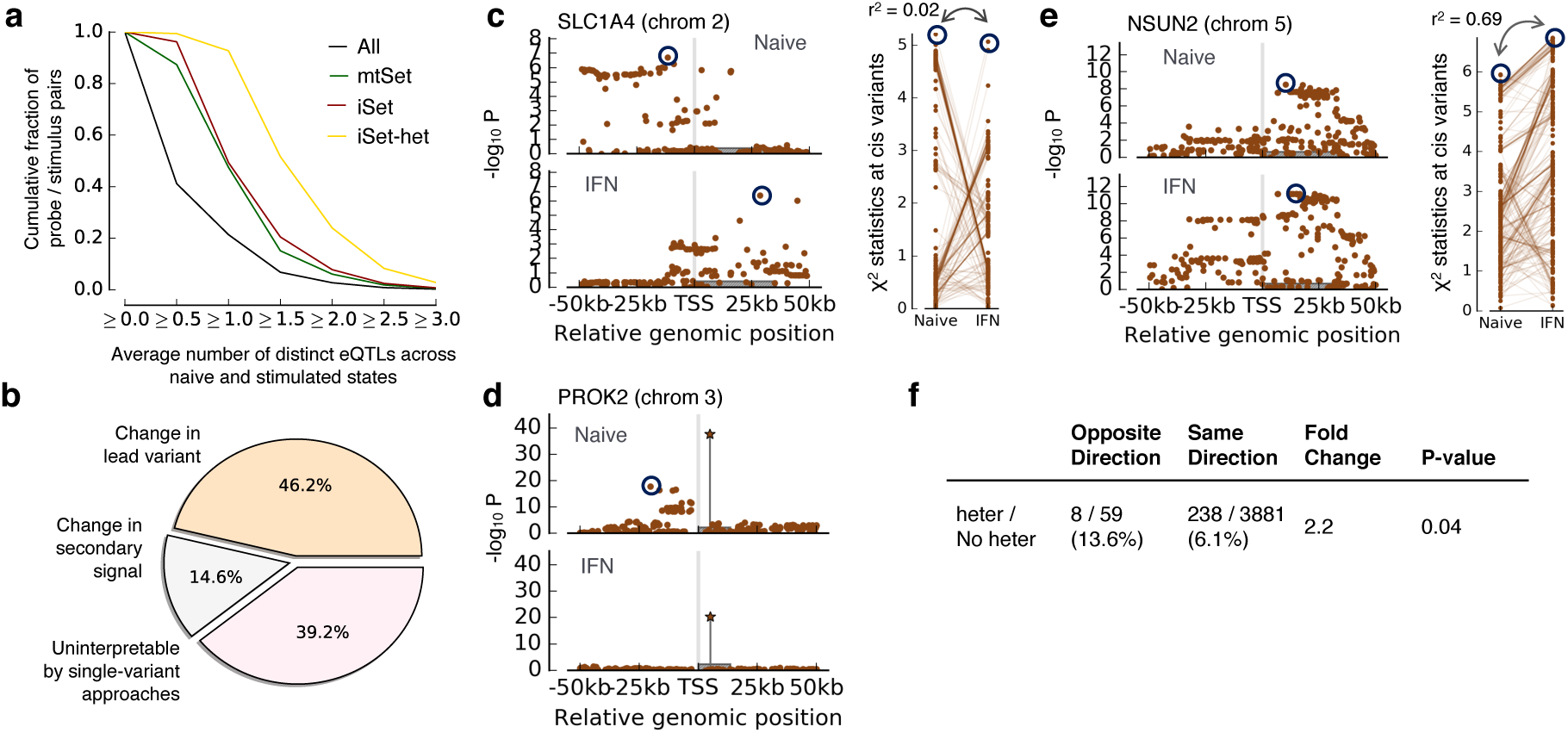
Characterization of genes with significant heterogeneity GxC for stimulus eQTLs in monocytes. **(a)** Cumulative fraction of probe/stimulus pairs with increasing numbers of distinct univariate eQTLs (average of the naïve and the stimulated state using step-wise selection) for different gene sets (**Methods**). Shown are cumulative fractions of all probe/stimulus pairs (All), those with significant *cis* associations (mtSet), pairs with significant GxC (iSet) and instances with significant heterogeneity GxC(iSet-het).**(b)** Breakdown of 1,281 probe/stim ulus pairs with significant hetero geneity GxC into distinct classes defined using the results of a single-variant step-wise LMM **(Methods)**. **(c-e)** Manhattan plots for representative probes with significant heterogeneity GxC effects. Grey boxes indicate the gene body. **(c)** Manhattan plot (left) and χ^2^ statistics for variants in both contexts (right) for the gene *SLC1A4*. Dark circles indicate distinct lead variants in both contexts (r^2^<0.2). **(d)** Manhattan plot after conditioning on the lead variant (secondary associations in the stepwise LMM) for the gene *PROK2*. The star symbol indicates the shared lead variant in both contexts. The conditional analysis reveals a secondary association that is specific to the naïve state. **(e)** Analogous plot as in c for the gene *NSUN2*, for which the single-variant model did not provide an interpretation of heterogeneity-GxC. **(f)**Breakdown of probe / stimulus pairs with shared lead variants, stratified by concordance of the effect direction (opposite-direction versus same-direction eQTLs) and significance of the heterogeneity-GxC test (heter vs No heter; FDR 5%). eQTLs with opposite effects were enriched for significant heterogeneity-GxC (2.2 fold enrichment, P<4e-2).

When overlaying heterogeneity GxC eQTLs detected using iSet-het with the results obtained from the single-variant step-wise LMM, we could attribute 46.2% of the heterogeneity-GxC effects (524 out of 1,135) to context-specific lead variants (defined using r^2^<0.2, FDR<5%, Fig. 4b,c, **Methods**). For an additional 14.6% of the heterogeneity eQTLs (166/1,135) the lead variants from a single-variant analysis were in high LD (r^2^>0.8), with context-specific secondary effects (Fig. 4b,d). The remaining 445 heterogeneity eQTLs (39.2%) could not be annotated using single-variant models.

One reason why heterogeneity GxC effects cannot be annotated using a single-variant model are differences in power. Indeed, for 22.2% of the heterogeneity-GxC cases without a single-variant interpretation (99/445), the single-variant LMM did not yield a significant effect in either of the two contexts (**Supplementary Fig. 13a**). For an additional 58.2% of the unannotated heterogeneity GxC effects (259/445), the single-variant LMM lead variants were in weak linkage (0.2<r^2^<0.8 example in Fig. 4e), which neither confirms nor rules out distinct genetic effects. One explanation for these instances are distinct polygenic architectures in both contexts. Consistent with this possibility, we observed that genetic effects captured by a polygenic model in both contexts (Best linear unbiased predictor, **Methods**) were markedly less correlated for probes with significant heterogeneity-GxC (**Supplementary Fig. 13b,c, Methods**).

Finally, we explored the relationship between probes with heterogeneity GxC and opposite effects as defined using conventional single-variant models. Following [21], we classified associations as opposite effects (lead variants r^2^>0.8 with opposite effect directions, **Methods**), yielding 67 eQTLs with reversed effect directions between contexts. iSet-het detected significant heterogeneity-GxC for 8 of these eQTLs, a 2.2 fold enrichment (P<5e-2) compared to eQTLs with consistent effect directions between contexts (238 gene/stimulus pairs with significant heterogeneity-GxC out of 4,119 eQTLs with consistent direction, Fig. 4f). Similar enrichments were also observed when considering individual stimulus contexts, resulting in significant enrichments for two out of three stimulus contexts (P<5e-2, fold change>4 in naïve/IFN and naïve/LPS-24h, **Supplementary Table 5**). Among the genes with significant heterogeneity-GxC are *OAS1*, *LMNA* and *PTK2B*, opposite-effect eQTLs that have been reported in the primary analysis of the same data [21] (**Supplementary Fig. 14**).

#### Using iSet to test for interaction effects in stratified populations

Thus far, we have considered settings with repeat measurements, where the same phenotype is measured in all individuals and contexts. Next, we considered applications of iSet to studies where individuals are phenotyped in only one context (**Supplementary Fig.2**, **Methods**). This is a common strategy in investigation of genotype-context interactions, where a population is stratified using a context variable.

We considered simulations analogous to those for complete designs (Fig. 2) to validate iSet for this design. We again confirmed statistical calibration of iSet (**Supplementary Fig. 15a**) and found similar power benefits as for complete designs (Fig. 5a,b, **Supplementary Fig. 15b,c**). In addition to single-variant LMMs, we also compared to a recently proposed set test for interactions (GESAT; [13]), which is designed for stratified populations. Notably, iSet was consistently better powered than GESAT, most likely because GESAT does not model correlations of the local genetic effect between contexts (**Methods, Supplementary methods**).

**Figure 5.**
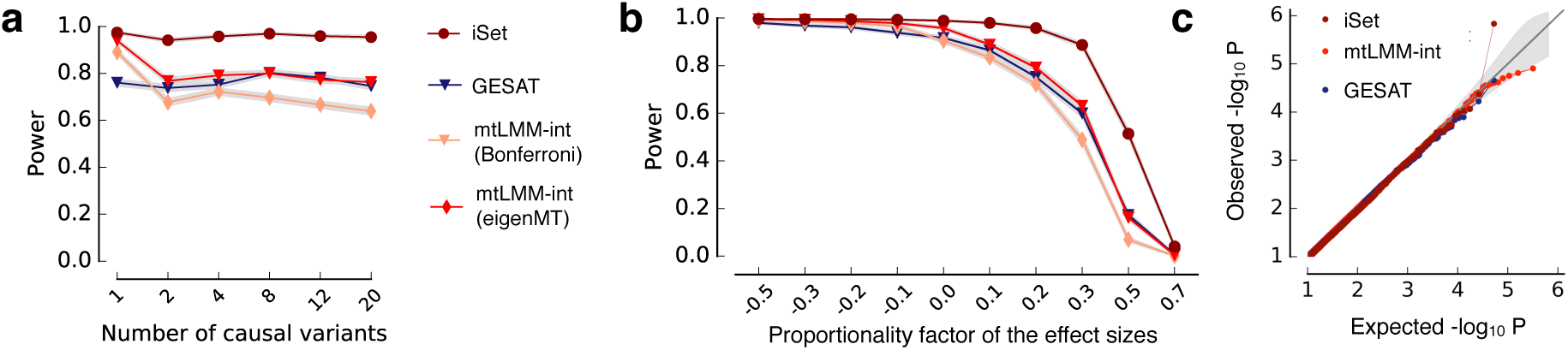
Application of iSet to stratified designs. **(a,b)** Power comparison of iSet and alternative methods using simulated data where each individual is phenotyped in one of two contexts. Shown is a comparison of power for alternative methods. **(a)** Power to detect interactions when simulating rescaling-GxC for increasing numbers of causal variants. **(b)** Power when varying the factor of proportionality of the variant effect sizes between contexts. Considered were iSet, a single-variant interaction test (mtLMM-int, [10]) as well as the interaction sequence kernel association test (GESAT, [13]), a set test designed for stratified populations. For single-variant models, two alternative approaches to adjust for multiple testing were considered (Bonferroni, eigenMT). **(c)** QQ-plot of P values from genotype-sex interaction tests for C-reactive protein levels using individuals from the Northern Finland Birth Cohort [20], considering the same methods.

Next, we applied iSet to test for genotype-sex interactions in four lipid-related traits (fasting HDL and LDL cholesterol levels, triglycerides and C-reactive protein) measured in 5,256 unrelated individuals from the Northern Finland Birth Cohort (NFBC1966 [20]). We tested consecutive 100kb regions (step size 50 kb; 52,819 genome-wide tests), and compared iSet to GESAT and the single-variant interaction test (**Methods**).

iSet retrieved one genome-wide significant interaction (C Reactive protein, chr1:40,450,000; P=1.47x10^−6^; FWER<10%), whereas alternative set tests and the single-variant models did not yield significant effects (Fig. 5c, **Supplementary Fig. 16,17, Supplementary Table 6**), even when using dense genotypes derived using imputation strategies (**Supplementary Fig. 18**). This interaction was replicated in a large meta-analysis (66,185 individuals [34]), which reports both an association for C-reactive protein at the same locus (P<6×10^-11^) as well as a marginal significant interaction with sex (P<5×10^-3^). Finally, a local single-variant analysis, separately for female and male individuals, provided evidence that this interaction reflects a male-specific genetic effect (**Supplementary Fig. 19**).

iSet revealed a second suggestive interaction with sex for LDL cholesterol levels (chr3:121,850,000, **Supplementary Fig. 16**). Although this effect failed genome-wide significance (FWER<20%), iSet again yielded stronger evidence than other methods (P_iSet_ = 3.7×10^-6^, P_GESAT_ = 4.8×10^-6^, P_mtLMM-int_ = 3.2×10^-5^). Among the genes at this locus is *ADCY5*, which has been linked to blood glucose levels in large meta-analyses [35, 36] and hence is a plausible candidate to affect LDL via glucose regulation [37].

Finally, we note that context stratification of quantitative traits can increase power for detecting associations rather than interactions, which is similar to previous strategies applied for single-variant analyses of quantitative [38] and categorical traits [39, 40]. Using this generalized association test, we identified three additional associations that were missed by conventional set tests and other methods (**Supplementary Fig. 16, Supplementary table 6**). These include the same locus with a sex-specific effect on C-reactive protein (chr1:40,450,000, P = 1.42×10^−7^ using mtSet, P=1.89 ×10^−3^ using a standard set test), and two associations for HDL cholesterol levels and triglycerides, both of which were replicated in larger meta analyses [41].

## Discussion

We have here proposed iSet, a method based on linear mixed models to test for gene-context interactions using variant sets. On simulated data as well as in applications to gene expression and human lipid-related traits, we have demonstrated that iSet yields increased power and improved interpretation for interaction effects compared to previous methods.

Methods for the joint analysis of multiple traits, including tests for genetic interactions, are not new per se. Most previous studies have used set-based methods to test for associations [7, 8, 12, 18], whereas tests for genotype-context interactions are still primarily carried out using single-variant models [10, 17]. iSet unifies several previous models (**Methods**), and uniquely offers set-based interaction tests on phenotypes in different contexts under the same or different (stratified) genetic backgrounds. Additionally, we have shown that set-based interaction tests can be useful to disentangle the genetic architecture of such loci, discerning consistent changes of genetic effects between contexts (rescaling-GxC) and changes in the configuration of causal variants (heterogeneity-GxC). The heterogeneity GxC test we propose is related to co-localization tests [19, 42, 43], however with a different objective.

In applications to a stimulus eQTL study we have shown that approximately 20% of the gene-stimulus interactions are associated with significant heterogeneity-GxC. This suggests that changes in the genetic architecture between stimulus contexts are relatively common. Additionally, we have observed that genes with opposite effects are enriched for heterogeneity-GxC. This finding points to a possible bias whereby opposite effects identified using single-variant models may in part be due to context-specific causal variants that are LD-tagged by a shared lead variant.

The proposed iSet model is not free of limitations. First, scalable inference in our model is achieved by exploiting the low-rank structure of variant sets, meaning that the number of variants in the analyzed region is typically small compared to the number of individuals. Similar to previous set-based tests [12], there are trade-offs between power and resolution, in particular when analyzing data from densely imputed or sequenced cohorts. General strategies for the design of optimal testing regions, for example using genome annotations and LD information, are an important area of future work. The inference scheme we have derived is efficient if phenotypes are either observed in all contexts and individuals or, alternatively, if a cohort is stratified using a context variable. Intermediate designs may also be considered, but currently require the use of separate imputation schemes [11, 44]. It is also worth noting that the test for heterogeneity-GxC (iSet-het) will be most accurate if all individuals are phenotyped in each context. Although in principle the model can also be used in stratified designs, there may be concerns that false positive heterogeneity GxC effects can arise due to technical factors, for example due to differences in genotyping accuracy or variant allele frequencies in the corresponding sub populations. A related issue is the need to choose the size of the region-set appropriately. While we find that the model is overall robust across a wide range of region sizes (**Supplementary Fig. 10**), the model will be best powered if the size of true causal regions approximately matches the testing region size, in particular for identifying heterogeneity-GxC effects.

Finally, we have here focused on pairwise analyses of different contexts. In principle the model could also be applied to analyze multiple related context and different traits, and the model could be extended to handle continuous environmental states, which currently require discretization. A related extension of the model is to test for genetic effects that are exclusive to one of the considered contexts. Developments in these directions are future work.

## Methods

### Software availability

iSet is freely available as part of the LIMIX package (http://github.com/PMBio/limix). Tutorials for using iSet either as command line tool or via a Python API can be found at https://github.com/PMBio/limix-tutorials/blob/master/iSet.

### The interaction set test (iSet)

To derive the model, we start assuming a fully observed design, where phenotypic measurements are available for all individuals and for each context. Briefly, the *N*×*C* phenotype matrix *Y* for *N* individuals and two or more contexts (*C*) is modeled as sum of fixed effects of *K* covariates, effects from *S* genetic variants in the region of interest (set component) and residual noise:

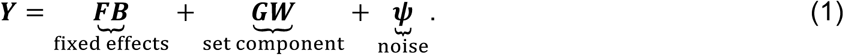

Here ***F*** (*N*×*K*) and ***G*** (*N*×*S*) denote respectively the fixed-effect covariates and the standardized genotypes of the variant set and ***B*** (*K*×*C*) and ***W*** (*S*×*C*) denote the corresponding effect sizes. The noise component ***ψ*** is assumed to follow a matrix-variate normal distribution, ***ψ***~MVN(0,***C****_n_*,***I****_N_*), where ***C****_n_* is a *C*×*C* covariance matrix that models residual covariances between traits. Note that in this formulation, population structure can be accounted for by including the leading principal component of the *N*×*N* (global) realized relatedness matrix [22] ?into the model as fixed effects [12]. In human populations, 10-20 principal components are typically sufficient to adjust for such structure [45]. Note that iSet can also account for population structure using an additional random effect term into the model (see **Supplementary Methods**). While computationally more expensive, this approach provides for additional robustness and calibration when analyzing cohorts with related individuals (see [12] for a discussion). All experiments reported here have been carried out using adjustment based on principal components, considering 10 PCs.

### Relationship between the trait-context covariance and the genetic architecture

To simplify the notation, we consider the case of two contexts, however the same derivation holds for larger numbers of contexts. Different genetic architectures between contexts are cast as specific assumptions on the *S*×*C* matrix of the variant effect sizes ***W***. A persistent genetic effect can be expressed as 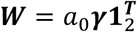, where a shared genetic signal ***γ*** (*S*×1) has a common scale *a*_0_ in both contexts. Rescaling-GxC can be expressed as *W* =***γa***^T^ where a common genetic signal ***γ*** (*S*×1) is modulated by context-specific scales *a* = [*a*_1_; *a*_2_] (2×1). Finally, in the most general case, the configuration of causal variants and their effects is independent between contexts, corresponding to *W* = [***γ***_1_,***γ***_2_]***AA^T^***, with genetic signals ***γ***_1_ (*S*×1) and ***γ***_2_ (*S*×1)and general scaling factor matrix ***A*** = [*a*_11_,*a*_12_,*a*_21_,*a*_22_] (2×2).

Marginalizing over the genetic signal ***γ***,***γ***_1_ and ***γ***_2_ assuming independent unit-variance normal prior distributions, results in a marginal likelihood of the form

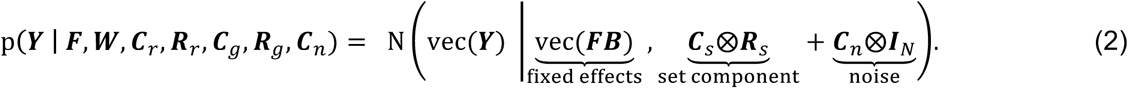

Here, vec denotes the stack-column operation, ⨂ the Kronecker product, ***C****_s_* is the *C×C* trait-context covariance for the set component and ***R****_S_* is the local realized relatedness matrix 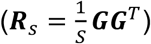. The stated alternative generative models for the structure of ***W*** have a one-to-one correspondence with alternative covariance structures for ***c****_S_* (**Supplementary Methods**), where for persistent effects***c****_S_* is a block covariance (***C****_s_* = *a*_0_**1**_2×2_), rescaling-GxC correspond to C_s_ with a rank-one structure (***c****_S_* = ***aa***^*T*^) and the most case of independent architectures in both contexts can be captured by a full rank covariance (***C****_s_* = ***AA***^*T*^). In order to test for associations, we additionally consider a null model without a set component (**C_s_ = 0**). The trait-context covariances ***c****_S_*,***c****_n_* and the fixed effect weights ***B*** are estimated using (restricted) maximum likelihood with constraints to obey the alternative structures of ***c****_S_*. Model parameters are optimized based on the restricted log marginal likelihood as objective, using a low-memory Broyden-Fletcher-Goldfarb-Shanno optizmier (L-BFGS) [46], implemented in the *fmin_l_bfgs_b* optimisation method of the SciPy python library. Specific tests are implemented as pairwise likelihood ratio (LLR) tests, considering LMMs with different trait-context covariances (See also Fig. 1b):

- **mtSet**: full-rank versus null
- **iSet**: full-rank versus block covariance
- **iSet**-het: full-rank versus rank-one

#### Obtaining P values

Empirical P values are estimated from the distribution of LLRs under the null. As permutation procedures are not well defined for interaction tests, for both iSet and iSet-het we generate test statistics from an empirical null distribution using a parametric bootstrap procedure [23]. Briefly, this procedure consists of sampling phenotypes from the null model with parameter values that maximize the likelihood on the observed data. Similarly to [8, 12], we consider a small number of parametric bootstraps for each region (typically 10-100 bootstraps) and pool the obtained null LLRs across all tested regions. The estimated distribution of null LLRs is used to obtain empirical P values. In an analysis of T genomic regions, the procedure to obtain P-values for the three tests can be summarized as follows:

- fit the no-association model (null), the block covariance model (block), the rank-one covariance model (rank-one) and the full-rank covariance model (full) and estimate LLRs for mtSet (full vs null), iSet (full vs block) and iSet-het (full vs rank-one);
- for each genomic region, sample J LLRs from the null for each of the three tests (we J permutations for mtSet, J parametric bootstraps for iSet and J parametric bootstraps for iSet-het);
- for each of the theses tests, pool the JT null LLRs across regions to obtain an empirical null and compute empirical P values.

Note that the number of parametric bootstraps / permutations will determine the minimum P value that can be obtained. For example, for T tests and B=30 bootstraps the minimum P value that can be estimated is 1/(BT), which correspond to a FWER of 1/B ≈ 0.03. While 30-100 bootstraps will be sufficient to reach typical thresholds in genome-wide studies, more stringent thresholds on significance levels (FWER<=1%) require a larger number of parametric bootstraps (see section below for computational considerations). For mtSet we use the same procedure but with permutations [12].

#### Data design, relatedness and scalability

Parameter inference using naïve implementations to fit the marginal likelihood model in iSet (Eqn. (2)) would scale cubically with the number of samples and contexts. iSet is optimized for cohorts with unrelated individuals, in which case population structure can be accounted for by including the top principal components (PCs) as fixed effect covariates. We have adapted prior work on set multi-trait tests, which for fully observed designs results in a computational complexity of 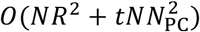, where *N* denotes the number of individuals, *R* denotes the number of variants in the region, *N*_PC_ is the number of PCs and *t* corresponds to the number of function evaluations of the optimizer (See [12] for details). iSet will be most efficient when the number of variants in the set is small compared to the number of individuals. To enable applications to stratified cohorts, we have extended this inference scheme to designs where phenotype data from each sample are observed in only one of the contexts, resulting in a computational complexity of 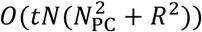 (**see Supplementary Methods**). For cohorts with related individuals and fully observed designs, iSet can also be applied with an additional random effect term in the model. In this case, we again re-use efficient inference schemes for multi-trait set tests [12]. This model is computationally more expensive, and scale with of *O*(*N*^3^ + *N*^2^*R* + *tNR*^2^). Note that for fully observed designs without the relatedness component, our implementation retains efficiency even if the number of genetic variants in the region set if greater than the number of individuals, i.e. it has complexity 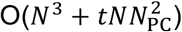. A tabular summary of the computational complexity of iSet for alternative data designs, confounder correction strategies and analysis settings is provided in **Supplementary Table 7**. Finally, while iSet is designed for the analysis of pairs of context, the model can also be applied to smaller numbers of contexts. In this case the method scales O(*tNC*^2^) with the number of contexts *C*.

Empirical runtime estimates in **Supplementary Fig. 1** were reported for different designs, using synthetic cohorts generated using data from the 1000 Genomes Project (phase 1, **Supplementary Methods**). We report the average per-region compute time measured on 100 regions with size 30 kb, considering a single core of an Intel Xeon CPU E5-2670 2.60-GHz to fit iSet. The runtime for all three considered tests, mtSet, iSet and iSet-het (including bootstraps) on the eQTL analysis took on average 28.7s per gene, resulting ~100h of compute time for a genome-wide analysis using a single core. Similarly, the runtime for all three tests for the NFBC data took was on average 107s per testing region, resulting in ~1,500h of compute time for a genome-wide analysis.

Finally, we note that the software implementation of iSet allows for efficient parallelization across multiple compute nodes and cores, similar to conventional GWAS approaches.

#### Variance decomposition model

The LMMs in iSet can also be used to estimate the phenotypic variance explained by the variant set for their persistent, rescaling-GxC and heterogeneity-GxC effects (**Supplementary Methods**).

#### Relationship to existing methods

iSet generalizes previous interaction set tests and multi-trait mixed models. Existing interaction set tests [13, 24–28] are designed for the analysis of stratified individuals and are not applicable to designs with repeat measurements, where the same trait is phenotyped in the same individuals in multiple contexts. Moreover, these existing methods do not account for correlated genetic effects within the region set and their underlying LMMs assume that the signal to noise ratio is identical in both contexts. The iSet model is more flexible and accounts for arbitrary genetic correlations and residuals covariances, using a null model that is similar to previous single-variant interaction tests [10]. iSet combines the advantages of several of these previous models; see **Supplementary Methods** for details.

#### Choice of the window size

As for any set test, the size of the region set is an important parameter in iSet. The specific choice will depend on the biological application, LD and marker density. We have previously explored trade-offs between the computational efficiency and power of association tests for different choices of the window size [12]. We here examined how the choice of the window size affects the power of detecting interaction and heterogeneity-GxC by considering sliding-window experiments with alternative windows sizes in simulations (**Supplementary Fig. 10**, see below). For the simulation experiments shown in Fig. 2, we considered sets with 30kb, which captures local LD in the data (**Supplementary Fig. 5**). For the analysis of the stimulus eQTL study, we have considered gene-based sets using a 100kb *cis* genetic region centered on the TSS, which is in line with other *cis* eQTL analyses [32]. Finally, for the genotype-sex interaction analysis in blood lipid levels we followed [12] and considered a sliding window approach with 100kb regions and a step size of 50kb.

#### Simulation study for fully observed designs

Simulations were carried out using a synthetic cohort of 1,000 individuals derived from genotypes of European populations in the 1000 Genomes project [29] (phase 1, 1,092 individuals, 379 Europeans). Following [12, 47], we composed synthetic genotypes as a mosaic of real genotypes from individuals of European ancestry, while preserving population structure (Supplementary Methods). We considered single-nucleotide polymorphism with a minor allele frequency of at least 2% (**Supplementary Fig. 4**). In all simulations, we simulated two contexts, modeled as the sum of a genetic contribution from a 30kb causal region, effects due to population structure, hidden covariates and identically distributed Gaussian noise. Effects due to population structure and hidden confounders were simulated with partial correlations across contexts, explaining variable proportions of the total phenotypic variance in each context (**Supplementary Table 2, Supplementary Methods**).

#### Statistical calibration

To assess the calibration of P values obtained from the interaction test (iSet) and the test for heterogeneity-GxC (iSet-het), we considered 100,000 datasets with two contexts where only persistent genetic effects (no interactions) were simulated (Fig. 2a). For each simulation we randomly selected a 30kb region and generated phenotypes simulating persistent effects from four causal variants and tested for GxC interaction in the region. To estimate P values, we used 30 parametric bootstraps for each test, resulting in a total of 3,000,000 null LLRs to estimate P values. Analogously, we assessed the calibration of iSet-het, where exclusively rescaling-GxC effects were simulated (**Supplementary Fig. 3**). Again, we considered 30 parametric bootstraps for each test and pooled LLRs to estimate P values.

#### Comparison with alternative methods

For comparison, we considered single-variant interaction tests as in [10] (mtLMM-SV-int), using an implementation in LIMIX [48]. To obtain region-based P values, we considered the minimum P value across all variants in the region, following adjustment for multiple testing. We consider two alternative strategies to adjust for multiple testing within variant sets: i) a conservative Bonferroni approach and ii) the recently proposed eigenMT model [30], which adjusts for the effective number of independent tests estimated based on the local LD structure. Existing set tests are not applicable for fully observed designs and hence were not considered (**Supplementary Methods**).

#### Power comparison

To assess power of iSet for alternative genetic architectures, we simulated interaction effects from a 30kb region either considering rescaling-GxC effects or more general effects that include heterogeneity-GxC, using the simulation settings in **Supplementary Table 2**. The total variance explained by the causal region across all traits was set to 2%. In the case of rescaling-GxC, we varied i) the number of causal variants in the region (from 1 to 20; Fig. 2b), and ii) the proportionality factor of the effect sizes between contexts (from −1 to 1, Fig. 2c). When simulating general effects that include heterogeneity-GxC, we randomly selected an equal number of context-specific causal variants and monitored the correlation of the total simulated genetic effects across contexts, thereby controlling the extent of heterogeneity-GxC. Again, local genetic effects were simulated to explain 2% of the total phenotypic variance in each context. We varied (see **Supplementary Table 2**) i) the extent of simulated heterogeneity-GxC (Fig. 2d) and ii) the total number of causal variants across contexts (**Supplementary Fig. 8**). For each parameter setting, we considered 1,000 repeat experiments. To obtain P values for set tests we considered 30 parametric bootstraps for each test and computed empirical P values from 30,000 null LLRs in each simulated scenarios. We used the Benjamini-Hochberg procedure to adjust for multiple testing across repeat experiments and assessed all methods in terms of power at a fixed FDR<5%.

#### Illustration case

For the simulated example region to illustrate iSet and alternative genetic architectures (Fig. 1c,d), we used a simulation procedure analogous to the strategy described above. Phenotypes were simulated as the sum of genetic effects from three distinct causal regions (30kb) within a 5Mb region on chromosome 13, harboring respectively persistent, rescaling-GxC and heterogeneity-GxC effects. The effects from individual regions was simulated to explain 5% of the total phenotype variance.

*Comparison of iSet-het with a baseline test for heterogeneity-GxC*

As an additional assessment of the accuracy of iSet-het to detect heterogeneity-GxC effects, we tested how well the model discriminates between regions with and without simulated heterogeneity-GxC. We considered the identical 10,000 regions in Fig. 2c for which no heterogeneity-GxC effects were simulated as well as the 10,000 regions in Fig. 2d with heterogeneity-GxC. We ranked all 20,000 regions based on the LLR of the heterogeneity test and used the receiver-operating characteristic (ROC) and precision-recall curves (**Supplementary Fig. 7**) to assess the ability of discriminating between these types of genetic effects. For comparison, we also considered a univariate baseline approach, scoring regions with significant associations using the squared Pearson correlation between the lead variants in both contexts (low squared Pearson correspond to high rank). We considered alternative significance thresholds on region-based P values obtained using eigenMT (P < 0.5, 0.01, 1e-2, 1e-3).

#### Choice of window size in sliding-window experiments

To study the effect of alternative sizes of the testing region on the power of iSet and iSet-het under different simulated scenarios, we considered sliding-window experiments using simulated data, when varying both the size of the simulated causal and of the testing region. Phenotypes were simulated across two contexts using the approach as described above, considering a causal region with variable size (30kb, 100kb, 300kb and 1Mb). For each of these simulation settings, we carried out a sliding window analysis in the surrounding 1 Mb region with testing windows of 30kb, 100kb, 300kb and 1Mb (the step size was set to the half of the size of the testing region). We used Bonferroni to adjust for multiple testing across regions. For comparison, we also considered the single-variant test for interactions (mtLMM-int), applied to the same variants in the 1Mb region, and used eigenMT to adjust for multiple testing across variants while accounting for LD. For each scenario, we considered 200 repeat experiments and assessed power at FDR=5%. We considered either simulated pure rescaling-GxC (**Supplementary Fig. 10a**) or more general effects (rescaling+heterogeneity-GxC, **Supplementary Fig. 10b-c)**. For both sets of simulations, we considered the default simulation parameter values (**Supplementary Table 2**).

## Monocyte eQTL dataset

### Data pre-processing

The dataset consists of gene expression levels from primary monocytes, both in a naïve state and three different stimulus contexts, profiled in 432 genotyped individuals of European ancestry. Gene expression levels in the naïve state, after exposure to IFN-γ, after 24-hour LPS and after 2-hour LPS were available for 414, 367, 322 and 261 individuals respectively. Normalization, correction for batch and probe filtering were done as in [21]. Following [21], we only considered probes that (i) map to only one genomic location, (ii) do not overlap with SNPs (MAF>1% in Europeans populations of 1000 Genomes Project), (iii) map to regions on autosomal chromosomes, and (iv) were detected in sufficient number of samples (see [21] for more details). Additionally, we discarded probes that could not be mapped to ensemble gene IDs. Collectively, these filters resulted in 12,677 probes for analysis (out of 15,421). We further limited our analysis to the set of 228 individuals for which gene expression levels were available in all the four (stimulus) contexts. To account for hidden covariates and confounding factors, we applied PEER [49] with default parameter values, fitting 30 hidden factors across all samples (individuals and stimulus states). PEER residuals for each gene and context were quantile-normalized to a standard normal distribution and used for all genetic analysis. Again, following the primary analysis [21], genotypes were imputed against the 1000 Genomes Project reference panel. After excluding variants with MAF<4%, variants with low imputation score (<0.9) and variants that deviate from the Hardy-Weinberg equilibrium (pv<10^-3^), we were left with 5,729,118 genome-wide variants (4,967,901 unique variants).

### eQTL mapping

Association and interaction tests were carried out considering 100 kb regions centered on the transcription start site of genes corresponding to individual probes (**Supplementary Fig. 4**). All tests were applied considering a pair-wise approach, jointly testing for eQTLs in the naive state and one of the stimulated states, considering set tests for association (mtSet), interaction (iSet) and heterogeneity-GxC (iSet-het). For comparison we also applied a single-variant tests using the same variants, testing for association (mtLMM) and stimulus interaction (mtLMM-int). For single-variant tests, we estimated gene-level significance using the P value of the lead *cis* variant (adjusted within *cis* regions using eigenMT, [30]). Empirical P values for iSet and iSet-het were estimated from 30 parametric bootstraps pergene and stimulus, combining all null LLRs across probes (resulting in 380,310 null LLRs per stimulus overall). Empirical P values for mtSet were obtained using the same permutation procedure as in [12]. Results from all methods were adjusted for multiple testing across probes using the Benjamini Hochberg procedure applied to each stimulus context separately. Reported results correspond to significant effects at genome-wide FDR < 5% (**Fig. 3a, Supplementary Table 4**).

### Best linear unbiased predictor from single-context set test

To illustrate the properties of the heterogeneity-GxC QTLs detected by iSet-het, we additionally considered univariate set tests in the same *cis* regions, however independently modeling each cellular context. At FDR<5% this analysis revealed 4,187, 4,786, 4,240 and 4,620 probes with an eQTL respectively in the naive, IFN-gamma, LPS2h and LPS24h states (**Supplementary Table 4**). To estimate the cis-genetic contribution to geneexpression in each context we calculated the Best Linear Unbiased Predictor (BLUP) from the model as 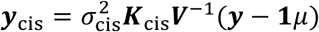, where *μ* is the estimated mean,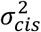 is the estimated variance explained by cis variants, ***K****_cis_* is the *cis* realized relatedness matrix, *V*^-1^ is the inverse of the total estimated covariance and *y* is the gene-expression vector in the corresponding context.

### Single-variant forward selection LMM

We used a single-variant forward selection LMM [33] to characterize eQTLs with significant heterogeneity-GxC effects. The model was fit considering up to three steps for gene and context, iteratively accounting for lead variant as additional fixed effect covariates when significant (FDR<5%). For each cellular context, region-based P values were adjusted for multiple testing across probes using the Benjamini Hochberg procedure for each of the three steps (only across probes that were tested at that specific step). This analysis yielded 15,756, 2,690 and 457 instances (across all genes and contexts) with one, two or three associations respectively (**Supplementary Table 4**).

Results from step-wise analysis were used to annotate probes with significant heterogeneity-GxC. We denoted the 1,449 probes that have significant marginal associations in both contexts and independent lead variants (*r*^2^ < 0.20) as a shift in lead variants between the two contexts. Probes with a shared lead eQTL (significant in both contexts, lead variants r^2^ > 0.80) were annotated using secondary effects. Among the 4,186 probes with shared main effects, this analysis revealed context-specific secondary QTLs were identified for 999 genes. Context-specific secondary effects were defined when either i) the secondary effect was significant in only one of the two contexts or ii) the secondary effects lead variants were in low LD (r^2^ < 0.20) (Fig. 4b).

### Annotation of opposite-effect eQTLs

We classified the 4,186 eQTLs with shared lead eQTL into directionally consistent and opposite-effect eQTLs. Briefly, opposite effects were defined by three criteria, i) marginal significance in both contexts, ii) LD between contexts (r2>0.8) and iii) negative correlation of genetic effects. These criteria resulted in 67 opposite-direction QTLs. Directionally consistent eQTLs correspond to criteria i) and ii) but positive correlated genetic effects, resulting in 4,119 co-located QTLs. Statistical significance of the enrichment for significant heterogeneity-GxC effects in opposite-direction eQTLs rather than same-direction eQTLs was assessed using a one-sided Fisher’s exact test (Fig. 4f).

### iSet for analysis of stratified cohorts

#### Simulations for analysis of stratified individuals

To study performance of iSet when considering interaction analyses in stratified cohorts, we considered simulation experiments analogous to those for fully observed designs. We generated a synthetic cohort of 2,000 Europeans where each individual was phenotyped in only in one of two contexts. For each individual, the phenotyped context was independently selected using a draw from a Bernoulli distribution (symmetric, 50% success rate). Statistical calibration and power simulations were performed analogously to the approach used for fully observed designs. Population structure was accounted for using the first ten principal components of the realized relatedness matrix as fixed effect covariates. We did not consider tests for heterogeneity-GxC, as differential tagging of causal variants could potentially result in spurious heterogeneity-GxC signals, and hence additional controls would be required. However, in principle the test applies to stratified populations.

#### Comparison to alternative methods

We compared iSet to the single-variant interaction tests as in [10] (mtLMM-int) and the gene-environment set association test (GESAT) [13]. The latter approach is representative for a family of closely related set tests that can only be applied to test for interaction effects in stratified populations (See **Supplementary Methods**). As an additional comparison, we extended the single-variant interaction test in [10] for stratified cohorts. To the best of our knowledge there are currently no implementations of mtLMM-int that can be applied to such designs. The models are available within the LIMIX package [48] (for full details see **Supplementary Methods**). GESAT was run using the function GESAT of the package iSKAT version 1.2. Both iSet and GESAT were applied on identically processed standardized variants.

#### Genotype-sex interaction tests in lipid traits

We performed a genotype-sex interaction analysis of four blood lipid phenotypes (C-reactive protein (CRP), triglycerides (TRIGL), LDL and HDL cholesterol levels) measured in 5,256 unrelated individuals from the NFBC1966 cohort [20] (phs000276.v1.p1). Following [11, 12], we regressed out major covariates, following a quantile-normalization of each trait individually. In order to correct for population structure, we considered the first ten principal components of the realized relatedness matrix as fixed effect covariates.

We applied mtSet and iSet to 318,653 genome-wide variants with an allele frequency of at least 1% using a sliding-window approach (100kb regions, 50kb step size; resulting in 52,819 windows overall; (**Supplementary Fig. 4**). For comparison we considered the single-variant interaction test [10], GESAT [13] and stSet [8], a univariate set test without stratification by sex. For each window we considered 100 permutations for mtSet and stSet and 100 parametric bootstraps for iSet and combined the obtained null LLRs across windows and traits (for a total of 21,127,600 null LLRs per test) to obtain empirical P values. Significance of the considered statistical tests was assessed at FWER=10%. Summary results from all considered methods are reported in **Supplementary Table 6.**

#### Imputation of NFBC1966 genotypes

Genotype data from NFBC1966.phs000276.v1.p1 were imputed using the 1000 Genomes Project phase 3 reference panel as described in the following. After aligning the dataset to the reference panel, we ran shapeit v2.r727 [50] with recommended parameters on each chromosome to produce haplotype estimates. We used impute2 v2.3.2 [51] with recommended parameters to impute untyped genotypes. Imputation was performed on chunks of approximately 5Mb. We merged region with less than 200 SNPs and avoided considering regions that span the centromere.

## Supplementary Information

### Tables

- Supplementary Table 1| Type-1 error estimates on simulated data.
- Supplementary Table 2| Simulation settings
- Supplementary Table 3| Relationship between the proportionality factor of the effect sizes used in simulations, fold change and relative direction of genetic effects across contexts.
- Supplementary Table 4 | Tabular summary of results from the monocyte gene expression analyses.
- Supplementary Table 5 | Enrichment analysis of heterogeneity-QTLs in opposite direction QTLs.
- Supplementary Table 6 | Tabular summary of the gene-by-sex interaction analysis in human blood lipid traits from NFBC1966 cohort.
- Supplementary Table 7| Computational complexity of iSet

Figures
- Supplementary Fig. 1 | Computational cost of iSet for alternative designs and cohort sizes.
- Supplementary Fig. 2 | Alternative study designs supported by iSet.
- Supplementary Fig. 3 | Statistical calibration of iSet-het when only rescaling-GxC effects are simulated.
- Supplementary Fig. 4 | Assessment of calibration under different types of model mismatch.
- SupplementaryFig. 5 | Distribution of the number of variants, the number of effective tests estimated by eigenMT and the average squared correlation within the testing regions in the different datasets.
- SupplementaryFig. 6 | Simulation results for synthetic genotypes without LD.
- SupplementaryFig. 7 | Comparison of iSet-het and single-variant strategies for discriminating rescaling from heterogeneity-GxC.
- SupplementaryFig. 8 | Power of iSet and iSet-het when simulating heterogeneity-GxC effects and increasing numbers of causal variants.
- SupplementaryFig. 9 | Assessment of genetic variance estimates from iSet using different covariance models.
- SupplementaryFig. 10 | Power of iSet and iSet-het in sliding window experiments with different sizes of the testing regions.
- Supplementary Fig. 11 | Number of positives for single-variant methods and set tests as a function of the false discovery rate in the monocyte stimulus QTL data.
- Supplementary Fig. 12 | Comparison of single-variant methods and set tests in the monocyte stimulus eQTL data.
- SupplementaryFig. 13 | Results from single-trait set tests applied to individual cellular contexts.
- SupplementaryFig. 14 | Examples of opposite-effect eQTLs with significant heterogeneity-GxC effects.
- Supplementary Fig. 15 | Calibration and power simulations for different GxE methods for analysis of stratified cohorts.
- Supplementary Fig. 16 | Manhattan plots from alternative methods applied for the genome-wide analysis of human lipid levels in the NFBC1966 cohort.
- Supplementary Fig. 17 | QQ plots when applying alternative methods to lipid levels in NFBC1966.
- Supplementary Fig.18 | Manhattan plot in the interaction locus for C-reactive protein using single-variant interaction tests on imputed variants.
- Supplementary Fig. 19 | The interaction for C-Reactive protein on chromosome 1 is a male-specific effect

## Acknowledgments

The authors would like to thank Na Cai, Chris Wallace, Leo Parts, Carl Anderson for comments on the manuscript. We thank Kaur Alasoo and the whole Stegle group for useful discussions. We thank Julian Knight and Benjamin Fairfax for sharing processed data for the monocyte stimulus eQTL study.

## References

1 Kang HM, Sul JH, Service SK, Zaitlen NA, Kong SY, Freimer NB, et al. Variance component model to account for sample structure in genome-wide association studies. Nature genetics. 2010;42(4):348–54. doi: 10.1038/ng.548. PubMed PMID: 20208533; PubMed Central PMCID: PMC3092069.

2 Yang J, Zaitlen NA, Goddard ME, Visscher PM, Price AL. Advantages and pitfalls in the application of mixed-model association methods. Nature genetics. 2014;46(2):100–6. doi: 10.1038/ng.2876. PubMed PMID: 24473328; PubMed Central PMCID: PMC3989144.

3 Rakitsch B, Stegle O. Modelling local gene networks increases power to detect transacting genetic effects on gene expression. Genome biology. 2016;17(1):33. doi: 10.1186/s13059-016-0895-2. PubMed PMID: 26911988; PubMed Central PMCID: PMC4765046.

4 Fusi N, Stegle O, Lawrence ND. Joint modelling of confounding factors and prominent genetic regulators provides increased accuracy in genetical genomics studies. PLoS computational biology. 2012;8(1):1002330. doi: 10.1371/journal.pcbi.1002330. PubMed PMID: 22241974; PubMed Central PMCID: PMC3252274.

5 Listgarten J, Kadie C, Schadt EE, Heckerman D. Correction for hidden confounders in the genetic analysis of gene expression. Proceedings of the National Academy of Sciences of the United States of America. 2010;107(38):16465–70. doi: 10.1073/pnas.1002425107. PubMed PMID: 20810919; PubMed Central PMCID:PMC2944732.

6 Wu MC, Kraft P, Epstein MP, Taylor DM, Chanock SJ, Hunter DJ, et al. Powerful SNP-set analysis for case-control genome-wide association studies. American journal of human genetics. 2010;86(6):929–42. doi: 10.1016/j.ajhg.2010.05.002. PubMed PMID: 20560208; PubMed Central PMCID: PMC3032061.

7 Chen H, Meigs JB, Dupuis J. Sequence kernel association test for quantitative traits in family samples. Genetic epidemiology. 2013;37(2):196–204. doi: 10.1002/gepi.21703. PubMed PMID: 23280576; PubMed Central PMCID: PMC3642218.

8 Listgarten J, Lippert C, Kang EY, Xiang J, Kadie CM, Heckerman D. A powerful and efficient set test for genetic markers that handles confounders. Bioinformatics. 2013;29(12):1526–33. doi: 10.1093/bioinformatics/btt177. PubMed PMID: 23599503; PubMed Central PMCID: PMC3673214.

9 Lippert C, Xiang J, Horta D, Widmer C, Kadie C, Heckerman D, et al. Greater power and computational efficiency for kernel-based association testing of sets of genetic variants. Bioinformatics. 2014;30(20):3206–14. doi: 10.1093/bioinformatics/btu504. PubMed PMID: 25075117.

10 Korte A, Vilhjalmsson BJ, Segura V, Platt A, Long Q, Nordborg M. A mixed-model approach for genome-wide association studies of correlated traits in structured populations. Nature genetics. 2012;44(9):1066–71. doi: 10.1038/ng.2376. PubMed PMID: 22902788; PubMed Central PMCID: PMC3432668.

11 Zhou X, Stephens M. Efficient multivariate linear mixed model algorithms for genome-wide association studies. Nature methods. 2014;11(4):407–9. doi: 10.1038/nmeth.2848. PubMed PMID: 24531419.

12 Casale FP, Rakitsch B, Lippert C, Stegle O. Efficient set tests for the genetic analysis of correlated traits. Nature methods. 2015;12(8):755–8. doi: 10.1038/nmeth.3439. PubMed PMID: 26076425.

13 Lin X, Lee S, Christiani DC, Lin X. Test for interactions between a genetic marker set and environment in generalized linear models. Biostatistics. 2013;14(4):667–81. doi: 10.1093/biostatistics/kxt006. PubMed PMID: 23462021; PubMed Central PMCID: PMC3769996.

14 Smith EN, Kruglyak L. Gene-environment interaction in yeast gene expression. PLoS biology. 2008;6(4):83. doi: 10.1371/journal.pbio.0060083. PubMed PMID: 18416601; PubMed Central PMCID: PMC2292755.

15 Anholt RR, Mackay TF. Quantitative genetic analyses of complex behaviours in Drosophila. Nature reviews Genetics. 2004;5(11):838–49. doi: 10.1038/nrg1472. PubMed PMID: 15520793.

16 Melchinger AE, Utz HF, Schon CC. Genetic expectations of quantitative trait loci main and interaction effects obtained with the triple testcross design and their relevance for the analysis of heterosis. Genetics. 2008;178(4):2265–74. doi: 10.1534/genetics.107.084871. PubMed PMID:18430948; PubMed Central PMCID:PMC2323814.

17 Sul JH, Bilow M, Yang WY, Kostem E, Furlotte N, He D, et al. Accounting for Population Structure in Gene-by-Environment Interactions in Genome-Wide Association Studies Using Mixed Models. PLoS genetics. 2016;12(3):1005849. doi: 10.1371/journal.pgen.1005849. PubMed PMID: 26943367; PubMed Central PMCID: PMC4778803.

18 Brown BC, Price AL, Patsopoulos NA, Zaitlen N. Local Joint Testing Improves Power and Identifies Hidden Heritability in Association Studies. Genetics. 2016. doi: 10.1534/genetics.116.188292. PubMed PMID: 27182951.

19 Wallace C. Statistical testing of shared genetic control for potentially related traits. Genetic epidemiology. 2013;37(8):802–13. doi: 10.1002/gepi.21765. PubMed PMID: 24227294; PubMed Central PMCID: PMC4158901.

20 Sabatti C, Service SK, Hartikainen AL, Pouta A, Ripatti S, Brodsky J, et al. Genome-wide association analysis of metabolic traits in a birth cohort from a founder population. Nature genetics. 2009;41(1):35–46. doi: 10.1038/ng.271. PubMed PMID: 19060910; PubMed Central PMCID: PMC2687077.

21 Fairfax BP, Humburg P, Makino S, Naranbhai V, Wong D, Lau E, et al. Innate immune activity conditions the effect of regulatory variants upon monocyte gene expression. Science. 2014;343(6175):1246949. doi: 10.1126/science.1246949. PubMed PMID: 24604202; PubMed Central PMCID: PMC4064786.

22 Lee SH, Goddard ME, Visscher PM, van der Werf JH. Using the realized relationship matrix to disentangle confounding factors for the estimation of genetic variance components of complex traits. Genetics, selection, evolution: GSE. 2010;42:22. doi: 10.1186/1297-9686-42-22. PubMed PMID: 20546624; PubMed Central PMCID: PMC2903499.

23 Buzkova P, Lumley T, Rice K. Permutation and parametric bootstrap tests for gene-gene and gene-environment interactions. Annals of human genetics. 2011;75(1):36–45. doi: 10.1111/j.1469-1809.2010.00572.x. PubMed PMID: 20384625; PubMed Central PMCID: PMC2904826.

24 Chatterjee N, Kalaylioglu Z, Moslehi R, Peters U, Wacholder S. Powerful multilocus tests of genetic association in the presence of gene-gene and gene-environment interactions. American journal of human genetics. 2006;79(6):1002–16. doi: 10.1086/509704. PubMed PMID: 17186459; PubMed Central PMCID: PMC1698705.

25 Tzeng JY, Zhang D, Pongpanich M, Smith C, McCarthy MI, Sale MM, et al. Studying gene and gene-environment effects of uncommon and common variants on continuous traits: a marker-set approach using gene-trait similarity regression. American journal of human genetics. 2011;89(2):277–88. doi: 10.1016/j.ajhg.2011.07.007. PubMed PMID: 21835306; PubMed Central PMCID: PMC3155192.

26 Lin X, Lee S, Wu MC, Wang C, Chen H, Li Z, et al. Test for rare variants by environment interactions in sequencing association studies. Biometrics. 2016;72(1):156–64. doi:10.1111/biom.12368. PubMed PMID: 26229047; PubMed Central PMCID: PMC4733434.

27 Chen H, Meigs JB, Dupuis J. Incorporating gene-environment interaction in testing for association with rare genetic variants. Human heredity. 2014;78(2):81–90. doi: 10.1159/000363347. PubMed PMID: 25060534; PubMed Central PMCID: PMC4169076.

28 Zhao G, Marceau R, Zhang D, Tzeng JY. Assessing gene-environment interactions for common and rare variants with binary traits using gene-trait similarity regression. Genetics. 2015;199(3):695–710. doi:10.1534/genetics.114.171686. PubMed PMID: 25585620; PubMed Central PMCID: PMC4349065.

29 Genomes Project C, Abecasis GR, Auton A, Brooks LD, DePristo MA, Durbin RM, et al. An integrated map of genetic variation from 1,092 human genomes. Nature. 2012;491(7422):56–65. doi: 10.1038/nature11632. PubMed PMID: 23128226; PubMed Central PMCID: PMC3498066.

30 Davis JR, Fresard L, Knowles DA, Pala M, Bustamante CD, Battle A, et al. An Efficient Multiple-Testing Adjustment for eQTL Studies that Accounts for Linkage Disequilibrium between Variants. American journal of human genetics. 2016;98(1):216–24. doi: 10.1016/j.ajhg.2015.11.021. PubMed PMID: 26749306; PubMed Central PMCID: PMC4716687.

31 Gagneur J, Stegle O, Zhu C, Jakob P, Tekkedil MM, Aiyar RS, et al. Genotype-environment interactions reveal causal pathways that mediate genetic effects on phenotype. PLoS genetics. 2013;9(9):1003803. doi: 10.1371/journal.pgen.1003803. PubMed PMID: 24068968; PubMed Central PMCID: PMC3778020.

32 Consortium GT. Human genomics. The Genotype-Tissue Expression (GTEx) pilot analysis: multitissue gene regulation in humans. Science. 2015;348(6235):648–60. doi: 10.1126/science.1262110. PubMed PMID: 25954001; PubMed Central PMCID: 10.1126/science.1262110. PubMed PMID:25954001; PubMed Central PMCID:PMC4547484.

33 Segura V, Vilhjalmsson BJ, Platt A, Korte A, Seren U, Long Q, et al. An efficient multi-locus mixed-model approach for genome-wide association studies in structured populations. Nature genetics. 2012;44(7):825–30. doi: 10.1038/ng.2314. PubMed PMID: 22706313; PubMed Central PMCID: PMC3386481.

34 Dehghan A, Dupuis J, Barbalic M, Bis JC, Eiriksdottir G, Lu C, et al. Meta-analysis of genome-wide association studies in >80 000 subjects identifies multiple loci for C-reactive protein levels. Circulation. 2011;123(7):731–8. doi: 10.1161/CIRCULATIONAHA.110.948570. PubMed PMID: 21300955; PubMed Central PMCID: PMC3147232.

35 Dupuis J, Langenberg C, Prokopenko I, Saxena R, Soranzo N, Jackson AU, et al. New genetic loci implicated in fasting glucose homeostasis and their impact on type 2 diabetes risk. Nature genetics. 2010;42(2):105–16. doi: 10.1038/ng.520. PubMed PMID: 20081858; PubMed Central PMCID: PMC3018764.

36 Saxena R, Hivert MF, Langenberg C, Tanaka T, Pankow JS, Vollenweider P, et al. Genetic variation in GIPR influences the glucose and insulin responses to an oral glucose challenge. Nature genetics. 2010;42(2):142–8. doi: 10.1038/ng.521. PubMed PMID: 20081857; PubMed Central PMCID: PMC2922003.

37 Otero P, Herrera E, Bonet B. Dual effect of glucose on LDL oxidation: dependence on vitamin E. Free radical biology & medicine. 2002;33(8):1133–40. PubMed PMID: 12374625.

38 Kim YK, Kim Y, Hwang MY, Shimokawa K, Won S, Kato N, et al. Identification of a genetic variant at 2q12.1 associated with blood pressure in East Asians by genome-wide scan including gene-environment interactions. BMC medical genetics. 2014;15:65. doi: 10.1186/1471-2350-15-65. PubMed PMID:24903457; PubMed Central PMCID:PMC4059884.

39 Gauderman WJ, Zhang P, Morrison JL, Lewinger JP. Finding novel genes by testing G x E interactions in a genome-wide association study. Genetic epidemiology. 2013;37(6):603–13. doi: 10.1002/gepi.21748. PubMed PMID: 23873611; PubMed Central PMCID: PMC4348012.

40 Morris AP, Lindgren CM, Zeggini E, Timpson NJ, Frayling TM, Hattersley AT, et al. A powerful approach to sub-phenotype analysis in population-based genetic association studies. Genetic epidemiology. 2010;34(4):335–43. doi: 10.1002/gepi.20486. PubMed PMID: 20039379; PubMed Central PMCID: PMC2964510.

41 Kraja AT, Vaidya D, Pankow JS, Goodarzi MO, Assimes TL, Kullo IJ, et al. A bivariate genome-wide approach to metabolic syndrome: STAMPEED consortium. Diabetes. 2011;60(4):1329–39. doi: 10.2337/db10-1011. PubMed PMID: 21386085; PubMed Central PMCID: PMC3064107.

42 Wallace C, Rotival M, Cooper JD, Rice CM, Yang JH, McNeill M, et al. Statistical colocalization of monocyte gene expression and genetic risk variants for type 1 diabetes. Human molecular genetics. 2012;21(12):2815–24. doi: 10.1093/hmg/dds098. PubMed PMID: 22403184; PubMed Central PMCID: PMC3363338.

43 Fortune MD, Guo H, Burren O, Schofield E, Walker NM, Ban M, et al. Statistical colocalization of genetic risk variants for related autoimmune diseases in the context of common controls. Nature genetics. 2015;47(7):839–46. doi: 10.1038/ng.3330. PubMed PMID: 26053495; PubMed Central PMCID: PMC4754941.

44 Dahl A, Iotchkova V, Baud A, Johansson A, Gyllensten U, Soranzo N, et al. A multiple-phenotype imputation method for genetic studies. Nature genetics. 2016;48(4):466–72. doi:10.1038/ng.3513. PubMed PMID: 26901065; PubMed Central PMCID:PMC4817234.

45 Wang K, Hu X, Peng Y. An analytical comparison of the principal component method and the mixed effects model for association studies in the presence of cryptic relatedness and population stratification. Human heredity. 2013;76(1):1–9. doi: 10.1159/000353345. PubMed PMID: 23921716.

46 Zhu C, Byrd RH, Lu P, Nocedal J. Algorithm 778: L-BFGS-B: Fortran subroutines for large-scale bound-constrained optimization. ACM Transactions on Mathematical Software (TOMS). 1997;23(4):550–60.

47 Loh PR, Tucker G, Bulik-Sullivan BK, Vilhjalmsson BJ, Finucane HK, Salem RM, et al. Efficient Bayesian mixed-model analysis increases association power in large cohorts. Nature genetics. 2015;47(3):284–90. doi: 10.1038/ng.3190. PubMed PMID: 25642633; PubMed Central PMCID: PMC4342297.

48 Lippert C, Casale FP, Rakitsch B, Stegle O. LIMIX: genetic analysis of multiple traits. bioRxiv. 2014.

49 Stegle O, Parts L, Piipari M, Winn J, Durbin R. Using probabilistic estimation of expression residuals (PEER) to obtain increased power and interpretability of gene expression analyses. Nature protocols. 2012;7(3):500–7. doi: 10.1038/nprot.2011.457. PubMed PMID: 22343431; PubMed Central PMCID: PMC3398141.

50 Delaneau O, Marchini J, Genomes Project C, Genomes Project C. Integrating sequence and array data to create an improved 1000 Genomes Project haplotype reference panel. Nature communications. 2014;5:3934. doi: 10.1038/ncomms4934. PubMed PMID: 25653097; PubMed Central PMCID: PMC4338501.

51 Howie B, Fuchsberger C, Stephens M, Marchini J, Abecasis GR. Fast and accurate genotype imputation in genome-wide association studies through pre-phasing. Nature genetics. 2012;44(8):955–9. doi: 10.1038/ng.2354. PubMed PMID: 22820512; PubMed Central PMCID: PMC3696580.

